# Spatial control of viscoelasticity in phototunable hyaluronic acid hydrogels

**DOI:** 10.1101/646778

**Authors:** Erica Hui, Kathryn I. Gimeno, Grant Guan, Steven R. Caliari

## Abstract

Viscoelasticity has emerged as a critical regulator of cell behavior. However, there is an unmet need to develop biomaterials where viscoelasticity can be spatiotemporally tuned to mimic the dynamic and heterogeneous nature of tissue microenvironments. Toward this objective, we developed a modular hyaluronic acid hydrogel system combining light-mediated covalent and supramolecular crosslinking to afford spatiotemporal control of network viscoelastic properties. Covalently crosslinked elastic hydrogels or viscoelastic hydrogels combining covalent and supramolecular interactions were fabricated to match healthy and fibrotic liver stiffness. LX-2 human hepatic stellate cells cultured on viscoelastic substrates displayed reduced spreading, less actin stress fiber organization, and lower myocardin-related transcription factor A (MRTF-A) nuclear localization compared to cells on elastic hydrogels. We further demonstrated the dynamic capabilities of our hydrogel system through photomediated secondary incorporation of either covalent or supramolecular crosslinks to modulate viscoelastic properties. We used photopatterning to create hydrogel models with well-controlled patterned regions of stiff elastic mechanics representing fibrotic tissue nodules surrounded by regions of soft viscoelastic hydrogel mimicking healthy tissue. Cells responded to the local mechanics of the patterned substrate with increased spreading in fibrosis-mimicking regions. Together, this work represents an important step forward toward the creation of hydrogel models with spatiotemporal control of both stiffness and viscoelastic cell-instructive cues.

## 1. Introduction

The interplay between cells and their surrounding extracellular matrix (ECM) plays a critical role in regulating development, wound healing, and disease progression [1–3]. Through mechanisms such as mechanotransduction, a process in which mechanical forces are converted into biochemical signals, cells are constantly probing and responding to their evolving microenvironment [4]. Cell-ECM interactions are especially important in pathologies such as fibrosis, a heterogeneous pathological scarring process that can lead to irreversible loss of tissue function and organ failure. During fibrosis progression, healthy tissue mechanics transition from softer and viscoelastic to stiffer and less viscous [5,6]. Moreover, fibrosis progresses in a heterogeneous manner, leading to microscale spatial heterogeneity in the form of patchy, stiff fibrotic nodules surrounded by areas of softer, less affected tissue where nodule size often directly correlates with the severity of fibrosis [7–9]. The presence of a stiff microenvironment can guide mechanotransduction by providing necessary biophysical cues for activation of resident cells into fibrosis-promoting myofibroblasts [10], and elevated stiffness alone has been shown to drive progression of both fibrosis [11] and cancer [12].

Hydrogels have become valuable model systems to better understand the contributions that matrix biophysical properties play in regulating cell behaviors through their ability to mimic salient properties of natural tissue, including soft tissue mechanics and high water content [13,14], and numerous systems have already investigated the influence of hydrogel mechanics on cell behavior [3,9,15–21]. In particular, many groups have shown a direct correlation between increasing hydrogel Young’s modulus (stiffness) and elevated cell spreading in two-dimensional (2D) cultures [10,22–25]. Although many studies have developed homogenous substrates to study cell-ECM interactions, healthy and especially diseased tissues are inherently heterogeneous. During pathologies such as fibrosis, changes in the physical environment have direct implications on cell mechanotransduction, where activated cell patches begin depositing excessive amounts of ECM proteins, resulting in nodules of nonfunctional scar tissue [7]. Therefore, it is necessary to develop methods to recapitulate tissue heterogeneity in hydrogel models. Recent work using light-based chemistries to spatially pattern elastic substrates has shown that cells will exhibit behavior correlating to their local mechanics such as increased spreading on stiffer areas [9,26,27].

While these findings are informative, they typically involve covalently-crosslinked hydrogels that primarily behave as elastic solids and do not display time-dependent tissue-relevant mechanical properties. The majority of native tissues exhibit viscoelastic behaviors including stress relaxation, which can occur through both external and cell-mediated forces exerted onto the matrix. For this reason, viscoelasticity has recently emerged as a critical parameter for probing cell behaviors and functions. Viscoelastic hydrogels have been developed using ionic [16,17], supramolecular [28], and dynamic covalent crosslinking [29] mechanisms. Viscoelastic hydrogels with stress relaxation properties similar to native tissues have been shown to affect cell spreading, focal adhesion organization, proliferation, and differentiation in comparison with elastic hydrogels [16–19,30,31]. This can be attributed in part to cell-mediated reorganization and/or relaxation of the energy-dissipative viscoelastic hydrogel network. Recent work from Charrier *et al*. [19] showed changes in the behavior of hepatic stellate cells, the primary cellular source of hepatic myofibroblasts, when cultured on viscoelastic hydrogels. Stellate cells displayed lower spread area and reduced expression of α-smooth muscle actin (α-SMA), a hallmark of myofibroblast activation, with increasing hydrogel loss modulus [19]. This study highlighted the importance of how hydrogel viscoelasticity can regulate disease-relevant cellular behaviors.

While the importance of incorporating viscoelasticity into hydrogel cellular microenvironments is clearly established, an approach to spatially control viscoelastic properties in a manner that mimics heterogeneous tissue has not been developed. The ability to pattern regions of hydrogel stiffness and/or viscoelasticity in a manner that captures both the dynamic stiffening that occurs during fibrosis progression and the overall heterogeneity of fibrotic tissue would help establish more robust disease models to study pathological cell behaviors. Here, we designed a phototunable viscoelastic hydrogel system where stiffness and viscoelasticity can be independently tuned through control of network covalent and supramolecular interactions. Using this modular approach, we developed photopatterned substrates where stiffness and viscoelasticity could be spatially controlled and investigated the role that matrix mechanical properties played in regulating cell behavior in an *in vitro* model of fibrosis.

## 2. Materials and Methods

### 2.1. NorHA synthesis

Norbornene-modified HA was synthesized similar to previous methods [32]. Briefly, sodium hyaluronate (Lifecore, 74 kDa) was reacted with Dowex 50W proton exchange resin, filtered, titrated to pH 7.05, frozen, and lyophilized to yield hyaluronic acid tert-butyl ammonium salt (HA-TBA). HA-TBA was then reacted with 5-norbornene-2-methylamine and benzotriazole-1-yloxytris-(dimethylamino)phosphonium hexafluorophosphate (BOP) in dimethylsulfoxide (DMSO) for 2 hours at 25°C. The reaction was quenched with cold water, dialyzed (molecular weight cutoff, MWCO: 6-8 kDa) for 5 days, filtered, dialyzed for 5 more days, frozen, and lyophilized. The degree of modification was 22% as determined by ^1^H NMR (500 MHz Varian Inova 500, **Figure S1**).

### 2.2. β-CD-HDA synthesis

The synthesis of β-cyclodextrin hexamethylene diamine (β-CD-HDA) followed the procedure outlined previously [33]. p-Toluenesulfonyl chloride (TosCl) was dissolved in acetonitrile and added dropwise to an aqueous β-cyclodextrin (CD) suspension (5:4 molar ratio of TosCl to CD) at 25°C. After 2 hours, the solution was cooled on ice and an aqueous NaOH solution was added dropwise (3.1:1 molar ratio of NaOH to CD). The solution was reacted for 30 minutes at 25°C before adding ammonium chloride to reach a pH of 8.5. The solution was cooled on ice, precipitated using cold water and acetone, and dried overnight. The CD-Tos product was then charged with hexamethylene diamine (HDA) (4 g/g CD-Tos) and dimethylformamide (DMF) (5 mL/g CD-Tos), and the reaction was carried out under nitrogen at 80°C for 12 hours before being precipitated with cold acetone (5 × 50 mL/g CD-Tos), washed with cold diethyl ether (3 × 100 mL), and dried. The degree of modification was 61% as determined by ^1^H NMR (**Figure S2**).

### 2.3. β-CD-HA synthesis

β-cyclodextrin modified hyaluronic acid (β-CD-HA) was prepared through coupling of β-CD-HDA to HA-TBA. A reaction containing HA-TBA, 6-(6-aminohexyl)amino-6-deoxy-β-cyclodextrin (β-CD-HDA), and BOP in DMSO was carried out at 25°C for 3 hours. The reaction was quenched with cold water, dialyzed for 5 days, filtered, dialyzed for 5 more days, frozen, and lyophilized. The degree of modification was 27% as determined by ^1^H NMR (**Figure S3**).

### 2.4. Peptide synthesis

Solid phase peptide synthesis was performed on a Gyros Protein Technologies Tribute peptide synthesizer. Three thiolated adamantane peptides (Ad-KKK**C**G, KKK**C**G, and DDD**C**G) were synthesized on either Rink Amide MBHA high-loaded (0.78 mmol/g) or Wang (1 mmol/g) resins using standard solid supported Fmoc-protected peptide synthesis. The resin was swelled with 20% (v/v) piperidine in DMF and the amino acids were activated using HBTU and 0.4 N-methyl morpholine in DMF (5:1 excess). Peptides were cleaved in a solution of 95% trifluoroacetic acid, 2.5% triisopropylsilane, and 2.5% H2O for 2-3 hours, precipitated in cold ethyl ether, and dried overnight. The peptide product was re-suspended in H_2_O, frozen, and lyophilized. Synthesis was confirmed by MALDI (**Figures S4 and S5**).

### 2.5. HA hydrogel fabrication

2D hydrogel thin films were made between untreated and thiolated coverslips (50 μL, 18 x 18 mm). Elastic NorHA hydrogels were fabricated using ultraviolet (UV)-light mediated thiol-ene addition. 2 and 6 wt% NorHA hydrogel precursor solutions containing 1 mM thiolated RGD (GCGYG<u>RGD</u>SPG, Genscript) and dithiothreitol (DTT) were photopolymerized (5 mW/cm^2^) in the presence of lithium acylphosphinate (LAP) photoinitiator for 2 minutes. Viscoelastic NorHA-CDHA hydrogels were fabricated by first mixing CD-HA with the thiolated adamantane peptide (1:1 ratio of CD to adamantane, Ad) to introduce Ad-CD guest-host interactions before mixing in RGD, DTT, and NorHA. The 2 and 6 wt% NorHA-CDHA precursor solutions were then photopolymerized (5 mW/cm^2^, 2 minutes) in the presence of LAP. Hydrogels were swelled in PBS overnight at 37°C before subsequent cell seeding procedures.

### 2.6. Rheological characterization

All rheological measurements were performed at 25°C on an Anton Paar MCR 302 rheometer using a cone-plate geometry (25 mm diameter, 0.5°, 25 μm gap). Rheological properties were tested using oscillatory time sweeps (1 Hz, 1% strain) with a 2 minute UV irradiation (5 mW/cm^2^), oscillatory frequency sweeps (0.001-10 Hz, 1% strain), and cyclic stress relaxation and recovery tests alternating between 0.1% and 5% strain (1 Hz).

### 2.7. Cell culture

Human hepatic stellate cells (LX-2s [34], Millipore Sigma) were used between passages 6-8 for all experiments. Culture media contained Dulbecco’s Modified Eagle Medium (DMEM) supplemented with 10 v/v% fetal bovine serum (FBS, Gibco) and 1 v/v% penicillin/streptomycin/amphotericin B (10,000 U/mL, 10,000 μg/mL, and 25 μg/mL respectively, Gibco). For cell seeding, swelled thin film hydrogels (18 x 18 mm) were sterilized using germicidal UV irradiation for 2 hours and incubated in culture media for at least 30 minutes prior to cell seeding. Cultures were treated with 5-10 μg/mL mitomycin C (Sigma-Aldrich) in serum-free media for 2 hours, washed thrice with PBS, and incubated in complete culture media for at least 1 hour prior to cell seeding. Cells were seeded atop hydrogels placed in untreated 6-well plates at a density of 2 x 10^4^ cells per hydrogel. For all experiments, media was replaced every 2-3 days for 7 day cultures.

### 2.8. Immunocytochemistry, imaging, and analysis

For immunostaining, cell-seeded hydrogels were fixed in 10% buffered formalin for 15 minutes, permeabilized in 0.1% Triton X-100 for 10 minutes, and blocked in 3% bovine serum albumin (BSA) in PBS for at least 1 hour at room temperature. Hydrogels were then incubated overnight at 4°C with primary antibodies against myocardin-related transcription factor A (MRTF-A, rabbit polyclonal anti-Mk11 antibody, 1:600, Abcam) and either α-smooth muscle actin (α-SMA, mouse monoclonal anti-α-SMA clone 1A4, 1:400, Sigma-Aldrich) or rhodamine phalloidin to visualize F-actin (1:600, Invitrogen). The hydrogels were washed 3 times in PBS and incubated with secondary antibodies (AlexaFluor 488 goat anti-rabbit IgG, 1:800; AlexaFluor 555 goat anti-mouse, 1:800) for 2 hours in the dark at room temperature. The hydrogels were then rinsed 3 times with PBS and stained with a DAPI nuclear stain (1:10000) for 1 minute before rinsing twice with 3% BSA. Stained hydrogels were stored in the dark at 4°C until imaging. Microscopy was performed on a Zeiss AxioObserver 7 inverted microscope. 2D hydrogels were covered with an 18 x 18 mm glass coverslip and inverted for imaging. Exposure time and other image settings for each respective channel were held constant while imaging. Cell spread area, cell shape index (CSI) and MRTF-A nuclear localization were determined using a CellProfiler (Broad Institute, Harvard/MIT) pipeline modified to include adaptive thresholding. CSI determines the circularity of the cell, where a line and a circle have values of 0 and 1, respectively, and was calculated using the formula:

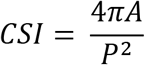

where *A* is the cell area and *P* is the cell perimeter. MRTF-A nuclear/cytosolic ratio was determined using the formula:

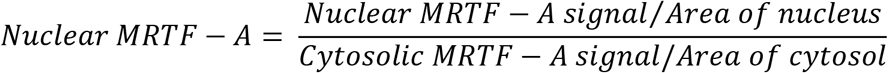

where the signal intensities were taken and normalized to their respective areas.

### 2.9. Photopatterning HA hydrogels

6 wt% NorHA hydrogels with low amounts of DTT (thiol:norbornene ratio = 0.12 for initially soft viscoelastic, 0.2 for initially soft elastic, 0.45 for initially stiff elastic) were fabricated and swelled overnight in PBS at 37°C. The hydrogels were first swelled in a 2 wt% BSA in PBS solution for 2 hours before being swelled in a 500 μL PBS solution containing 1 wt% BSA, LAP, DTT, and fluorescent thiolated peptide for 1 hour at 37°C before irradiation with a patterned photomask transparency (CAD/Art Services, Inc) for 2 minutes (5 mW/cm^2^). The resulting patterned hydrogels were washed with PBS several times prior to imaging or cell seeding.

### 2.10. Atomic force microscopy (AFM) characterization and analysis

AFM force spectroscopy was performed using an Asylum Research MFP 3D AFM. A silicon nitride cantilever (MLCT-O10/Tipless/Ti-Au, cantilever C, Bruker) with a nominal spring constant of 0.1 N/m was functionalized with a 25 μm diameter polystyrene bead at the tip. The spring constant of the cantilever was calibrated via thermal resonance curves prior to data collection. Nanoindentation tests were performed on photopatterned NorHA hydrogels in PBS to determine the mechanics of the patterned and non-patterned regions. Force versus distance curves were generated and the Young’s modulus at each indentation was calculated using the loading portion of the indentation curve by applying the Hertzian contact mechanics model and assuming a Poisson’s ratio of 0.5. Force relaxation tests were performed to study viscoelasticity of patterned substrates. Following indentation, the tip was held at a constant indentation depth for 10-30 seconds at a 500 Hz sampling rate. Indentation force and depth were recorded as a function of time [35].

### 2.11. Statistical analysis

Student’s t-tests (two experimental groups) or one-way ANOVA with Tukey’s HSD post hoc tests (more than two experimental groups) were performed for all quantitative data sets. All experiments included at least 3 hydrogels and/or 20 individual cells quantified per experimental group. Box plots of single cell data had error bars that were the lower value of either 1.5*interquartile range or the maximum/minimum value, with data points between 1.5*interquartile range and the maximum/minimum indicated as open circles. Significance was indicated by *, **, or *** corresponding to *P* < 0.05, 0.01, or 0.001 respectively.

## 3. Results and Discussion

### 3.1. Viscoelastic hydrogels were synthesized with a combination of covalent and supramolecular crosslinks

Hyaluronic acid was functionalized with norbornene groups (NorHA) to produce hydrogels containing a high degree of reactive sites (~ 20% of repeat units). Compared to common functional groups such as (meth)acrylates, which can react with each other to form kinetic chains, norbornene groups have high reactivity to thiyl radicals and low reactivity to themselves, allowing rapid and controllable thiol-ene click addition of both pendant and multifunctional thiolated groups [32]. This biorthogonal system was also chosen for its ability to easily synthesize hydrogels with a wide range of tissue-relevant mechanics by simple tuning of parameters such as crosslinker concentration or light intensity. In this study, soft (G’ ~ 0.5 kPa) and stiff (G’ ~ 5 kPa) thin film hydrogels were fabricated to represent healthy and fibrotic liver tissue, respectively [10,11]. Di-thiol molecules (DTT) were used to provide stable hydrogel crosslinks and thiolated RGD peptide (GCGYG<u>RGD</u>SPG) was incorporated to allow for cell attachment. Elastic NorHA hydrogels were fabricated via ultraviolet (UV)-light mediated thiol-ene addition between norbornenes on HA and thiols on DTT to create stable covalently-crosslinked networks.

Viscoelasticity was introduced to the system by incorporating reversible guest-host interactions between adamantane (guest) and β-cyclodextrin (host) groups. The adamantane (Ad) guest moiety has a high affinity to the hydrophobic cavity of β-cyclodextrin (*K*_*a*_ ~ 10^5^ M^−1^) and has previously been exploited to make viscoelastic, shear-thinning hydrogels [33,36,37]. For the viscoelastic hydrogel groups, β-cyclodextrin HA (CD-HA) and thiolated Ad peptide were mixed in solution (1:1 molar ratio of CD to Ad) to introduce supramolecular guest-host interactions, followed by the addition of NorHA and DTT (**Figure 1**). This particular methodology involving Ad peptide allowed for a more modular approach to fabricating hydrogels due to its detachment from the HA backbone prior to the thiol-ene addition, making the hydrogel precursors less viscous and easier to pipet and mix. Following mixing of the Ad peptide, CD-HA, NorHA, and DTT, the thiols on the cysteine residues of the CD-associated Ad peptide reacted with the norbornenes to form stable supramolecular connections between HA chains while the DTT formed covalent crosslinks, creating a viscoelastic hydrogel network with both covalent and supramolecular crosslinks.

**Figure 1.**
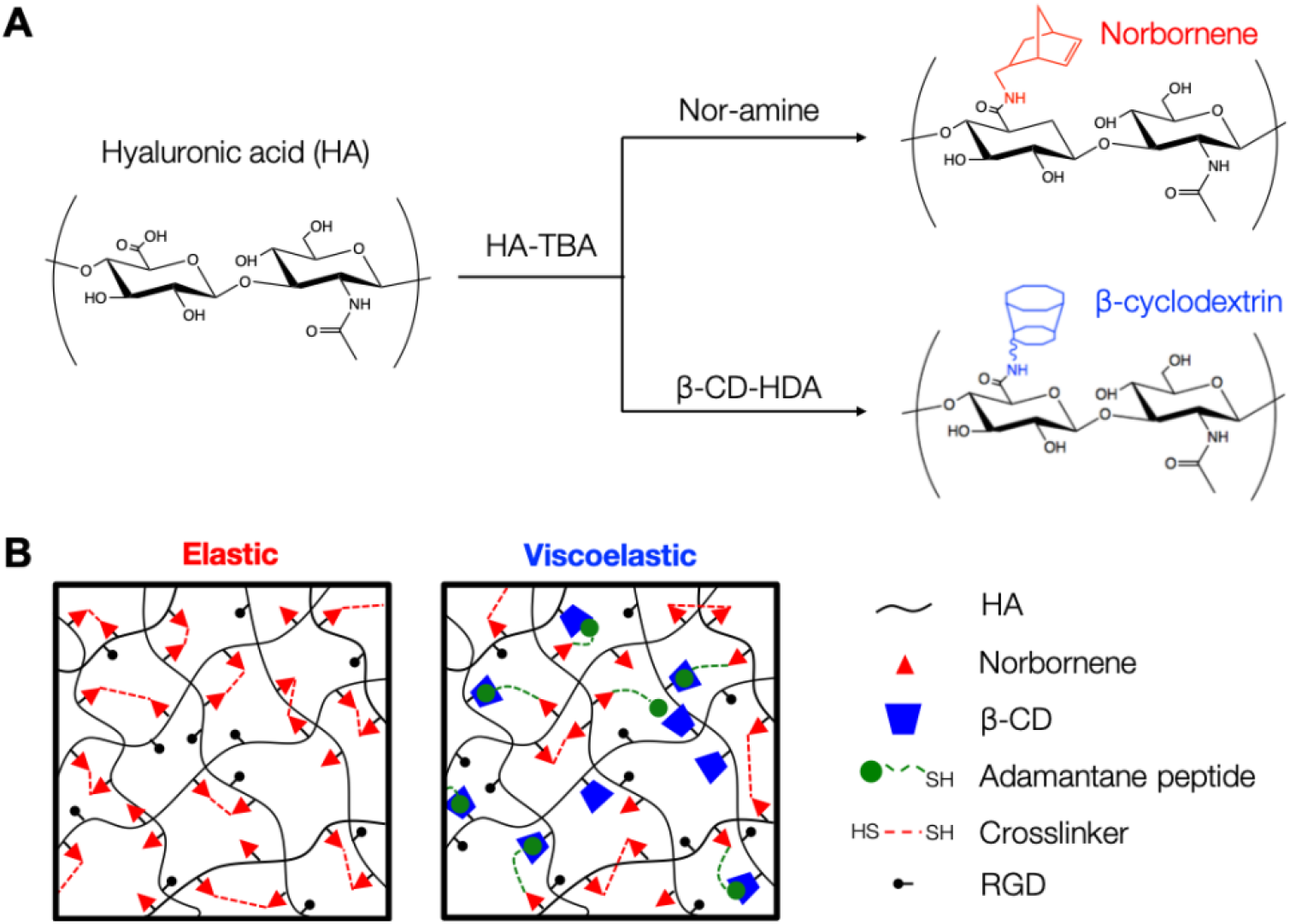
Overview of hydrogel synthesis and crosslinking. (A) Hyaluronic acid was first converted to HA-TBA salt before modification with norbornene or β-cyclodextrin groups using BOP coupling chemistry to synthesize NorHA and CD-HA. (B) For the elastic hydrogel system, covalent crosslinks between the norbornene groups were introduced using di-thiol crosslinkers via light-mediated thiol-ene addition. For the viscoelastic hydrogel system, thiol-ene photochemistry was used to introduce both stable supramolecular interactions between CD-HA and Ad groups on thiolated peptides in addition to di-thiol-mediated covalent crosslinks between the norbornenes.

### 3.2. Viscoelastic hydrogels display stress relaxation and frequency-dependent behavior

Hydrogel mechanical properties were characterized through shear oscillatory rheology (**Figure 2**). *In situ* gelation of hydrogel precursor solutions demonstrated rapid gelation kinetics controlled by light exposure, resulting in a nearly immediate plateau in storage and loss moduli once light irradiation was stopped (**Figures 2A and 2B**). Similar storage moduli were observed for the soft (elastic: G’ = 0.51 ± 0.08 kPa, viscoelastic: G’ = 0.46 ± 0.07 kPa) and stiff (elastic: 4.59 ± 0.24 kPa, viscoelastic: G’ = 4.93 ± 0.77 kPa) hydrogel groups corresponding to healthy and fibrotic liver tissue respectively. However, as expected, the viscoelastic hydrogels had significantly higher loss moduli (soft viscoelastic: G” = 67.2 ± 2.44 Pa, stiff viscoelastic: G” = 330 ± 45.5 Pa) compared to elastic groups (soft elastic: G” = 0.99 ± 0.89 Pa, stiff elastic: G” = 2.78 ± 2.21 Pa). Notably, the G” values for the viscoelastic hydrogels were within an order of magnitude of their G’, similar to the ratios observed in native viscoelastic tissue [19,38]. Hydrogel frequency sweeps revealed relatively constant storage and loss moduli for the elastic groups (**Figure 2C**). However, the viscoelastic hydrogels showed frequency-dependent behavior; at higher frequencies, the loss modulus increased, demonstrating that guest-host interactions were being disrupted with less time to re-associate. Stress relaxation and recovery tests showed that at a constant applied strain of 5%, the elastic hydrogels showed no stress relaxation over time due to their stable covalently-crosslinked network (**Figure 2D**). In contrast, the viscoelastic hydrogel groups showed cyclic stress relaxation in which high stress was observed, followed by a plateau to a final stress value equal to the corresponding elastic groups. The ability for the hydrogels to fully recover their mechanical properties upon repeated bouts of applied strain highlighted their viscoelasticity as opposed to viscoplasticity.

**Figure 2.**
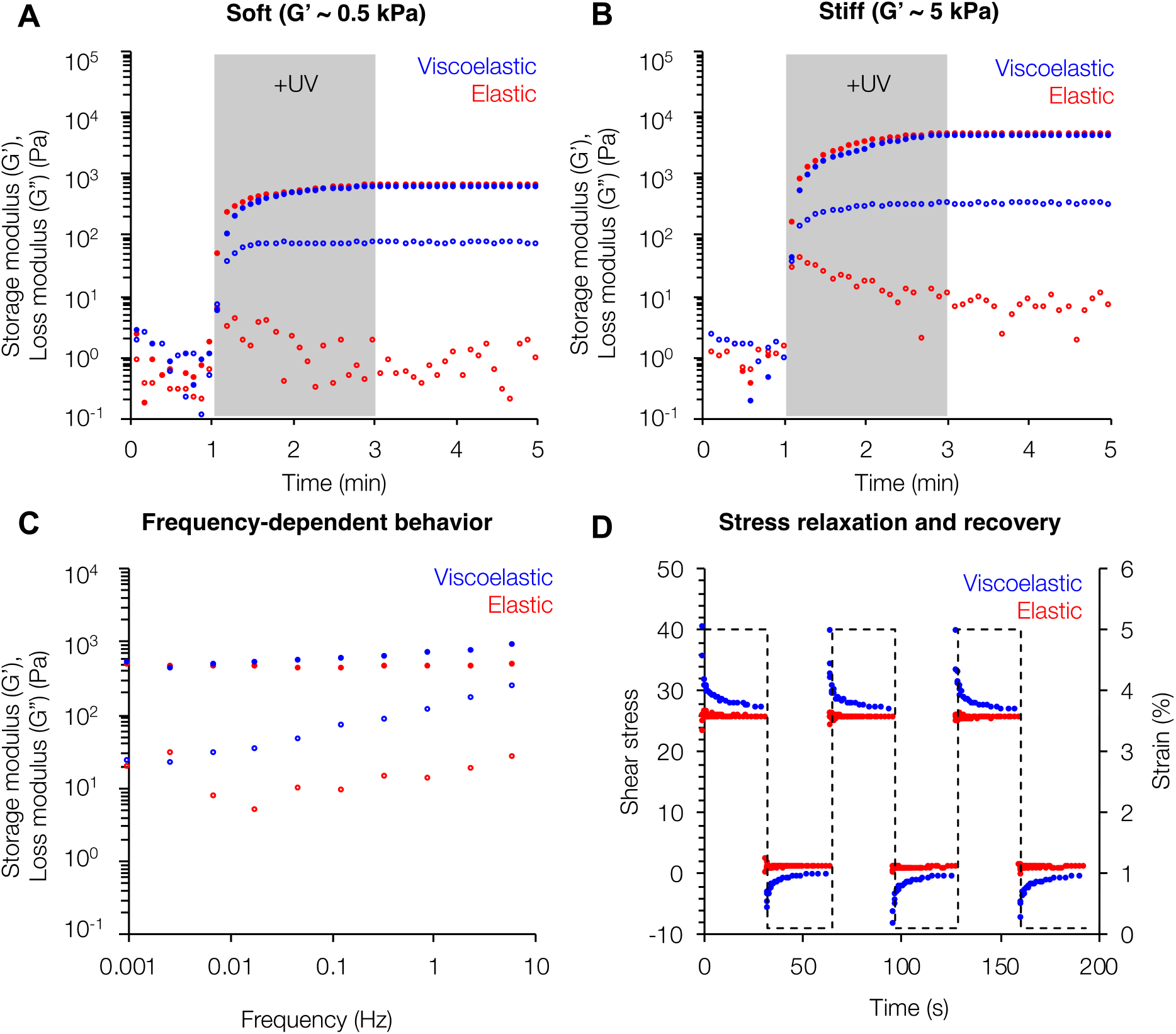
Rheological characterization of viscoelastic hydrogels. Viscoelastic hydrogels (*blue*) of equivalent storage moduli (*closed circles*) to their elastic counterparts (*red*) showed loss moduli (*open circles*) within an order of magnitude for both (A) “soft” (G’ ~ 0.5 kPa) and (B) “stiff” (G’ ~ 5 kPa) hydrogel formulations corresponding to healthy and fibrotic tissue respectively. (C) Viscoelastic hydrogels also showed frequency-dependent behavior with increasing loss moduli as frequency was increased, whereas the elastic hydrogel properties remained relatively constant. (D) Stress relaxation and recovery tests showed the full recovery of the mechanical properties of the viscoelastic hydrogels. For the frequency and stress relaxation tests, the soft groups are shown; similar trends were seen for the stiff groups (Figure S6).

### 3.3. Cell spreading is modulated by both hydrogel stiffness and viscoelasticity

After rheological characterization highlighted the tunable viscoelastic nature of our hydrogel design, we investigated the behavior of LX-2s, a human hepatic stellate cell line, in response to four hydrogel groups: soft elastic, stiff elastic, soft viscoelastic, and stiff viscoelastic. Cells on stiff elastic substrates showed increased spreading compared to cells on soft elastic substrates, similar to what has previously been reported for elastic substrates of increasing stiffness (**Figure 3**). In comparison the viscoelastic hydrogels, which had the same storage moduli as the corresponding elastic groups but higher loss moduli, supported decreased cell spreading and more rounded morphologies as measured by cell shape index (CSI) compared to their corresponding elastic substrates for both the soft and stiff groups. The differences in cell spreading and circularity were the greatest between cells cultured on the stiff elastic hydrogels, which became more elongated and extended protrusions (average spread area: 6623 μm^2^, CSI: 0.17), and cells cultured on the soft viscoelastic hydrogels, which showed smaller, more rounded morphologies (average spread area: 2982 μm^2^, CSI: 0.26). The reduction in stellate cell spreading is similar to results from a recent study where stellate cells showed reduced spreading and reduced expression of α-smooth muscle actin (α-SMA), a marker of myofibroblast activation, when cultured on polyacrylamide substrates with higher loss moduli [19]. Similarly, while cells on our viscoelastic hydrogels showed positive α-SMA staining, we also observed reduction in the organization of α-SMA stress fibers that is typical of activated myofibroblasts [19,39,40]. While around 85% of cells on stiff elastic hydrogels displayed at least some α-SMA stress fiber organization, only 6% and 22% of cells on soft and stiff viscoelastic hydrogels respectively displayed organized α-SMA stress fibers (**Figure S7**).

**Figure 3.**
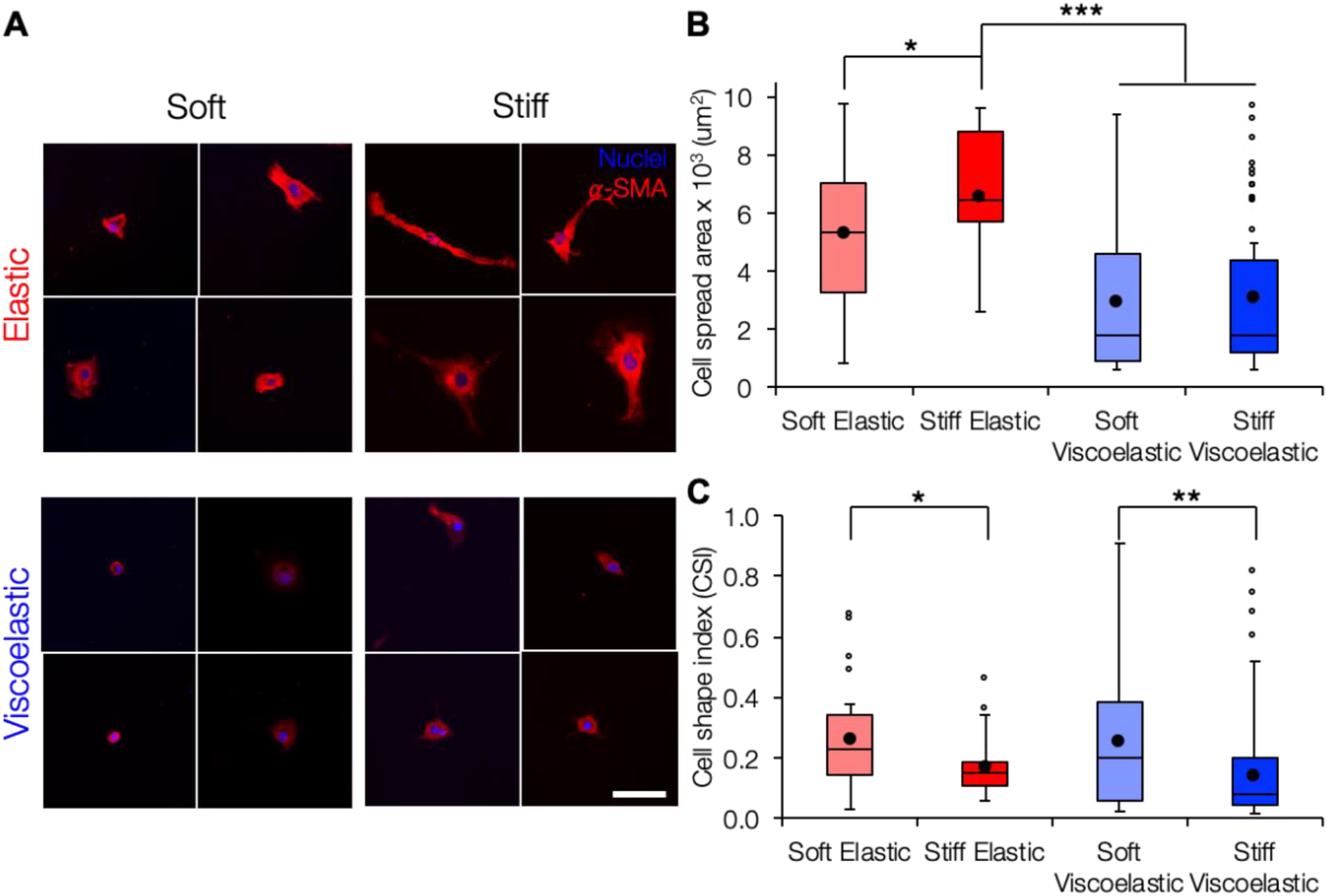
Cell spreading is modulated by both stiffness and viscoelasticity. (A) Representative images of LX-2 hepatic stellate cells stained for α-SMA (*red*) and nuclei (*blue*). Scale bar 100 μm. (B) While cell spreading was increased on stiff elastic compared to soft elastic hydrogels, spreading was significantly reduced on both soft and stiff viscoelastic hydrogels compared to the stiff elastic group. (C) Cell shape index was significantly higher for cells on soft hydrogels compared to their respective stiff counterparts, indicating that the cells displayed more rounded morphologies. *: *P* < 0.05, **: *P* < 0.01, ***: *P* < 0.001.

Since α-SMA expression is a relatively late marker of myofibroblast activation, we also sought to investigate earlier markers of fibrogenic mechanotransduction. Myocardin-related transcription factor A (MRTF-A), a transcriptional co-activator implicated in the regulation and progression of fibrosis, has been shown to drive α-SMA expression and subsequent myofibroblast activation [41–43]. Specifically, activation of mechanotransduction pathways through cell-matrix interactions can promote RhoA/ROCK signaling, actin polymerization, and subsequent MRTF-A nuclear translocation. MRTF-A then interacts with serum response factor (SRF), the transcription factor that promotes upregulation of the *Acta2* gene encoding for α-SMA [42,44–46]. We measured the ratio of MRTF-A nuclear to cytosolic signaling intensity and found elevated MRTF-A nuclear localization for cells on stiff compared to soft elastic hydrogels (**Figure S8**). However, cells cultured on the viscoelastic hydrogel groups showed reduced MRTF-A nuclear localization compared to the stiff elastic group. Overall, both soft and stiff viscoelastic hydrogels promoted reduced stellate cell spreading, α-SMA stress fiber organization, and MRTF-A nuclear localization. A possible explanation for these results could be the viscous dissipation of cell-generated forces into the matrix prevents spreading and activation of the mechanoresponsive signaling pathways investigated here [47].

### 3.4. Light-mediated thiol-ene addition enables secondary incorporation of covalent or supramolecular crosslinks

We demonstrated the dynamic capabilities of our viscoelastic hydrogel system through specific secondary introduction of either covalent or supramolecular interactions. First, we fabricated elastic hydrogels with increasing covalent crosslinking density controlled by sequential bouts of light exposure, permitting further thiol-ene crosslinking. Rheological analysis indicated that each additional irradiation corresponded with increasing storage modulus but relatively little change in loss modulus as expected for an elastic network (**Figure 4A**). Next, we made initially soft viscoelastic hydrogels containing unreacted norbornene and β-cyclodextrin groups and introduced additional supramolecular crosslinks through sequential thiol-ene addition of thiolated adamantane peptide. Each additional light irradiation led to increases in both the storage and loss moduli as the hydrogel maintained its viscoelastic nature (**Figure 4B**). Overall, the unique amenability of our system to the light-mediated introduction of either new covalent or supramolecular crosslinks sets the stage for creation of dynamic, heterogeneous viscoelastic hydrogels.

**Figure 4.**
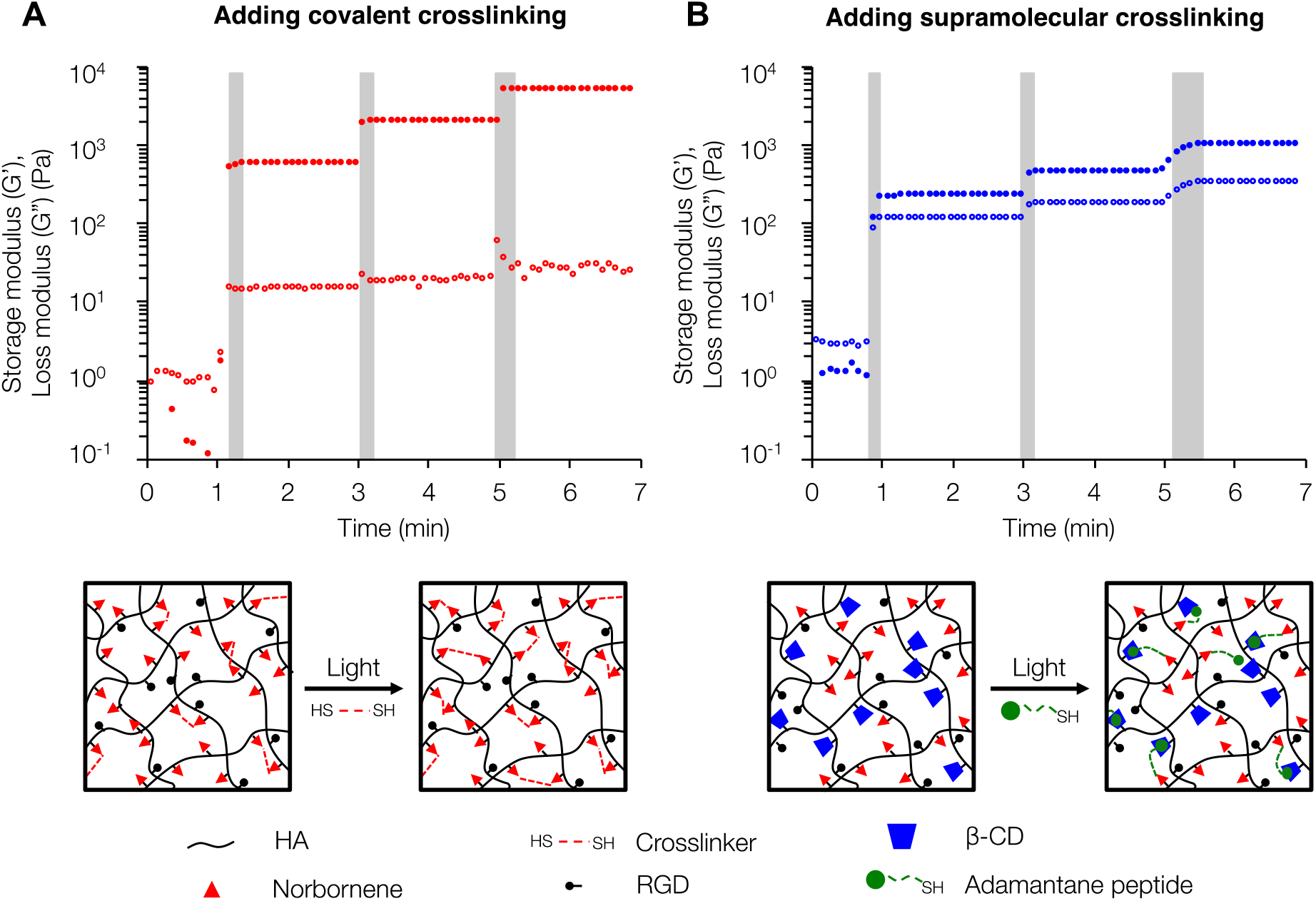
Secondary introduction of covalent or supramolecular crosslinks to modulate viscoelastic properties. (A) When incorporating new covalent crosslinks through DTT addition, each subsequent UV exposure (*gray bars*) results in an increased storage modulus but minor changes in loss modulus. (B) Following initial formation of a viscoelastic hydrogel, incorporating new supramolecular guest-host crosslinks leads to increases in both the storage and loss moduli with each UV exposure.

### 3.5. Photopatterning enables presentation of dynamic, heterogeneous, and cell-instructive viscoelastic hydrogel properties

After developing our viscoelastic hydrogel system and demonstrating its amenability to secondary crosslinking reactions, we explored the use of photopatterning to recapitulate the heterogeneity of matrix mechanical properties during fibrogenesis in a well-defined manner (**Figure 5**). Using photomasks to control light penetration into the hydrogel during secondary crosslinking enables spatial control over the thiol-ene addition reactions. Soft NorHA hydrogels were swelled in a solution containing 1 wt% BSA, LAP photoinitiator, DTT crosslinker or adamantane peptide, and thiolated fluorescent peptide. Fluorescence microscopy confirmed pattern fidelity, with alternating fluorescent and non-fluorescent regions present in the hydrogel (**Figure 5B**).

**Figure 5.**
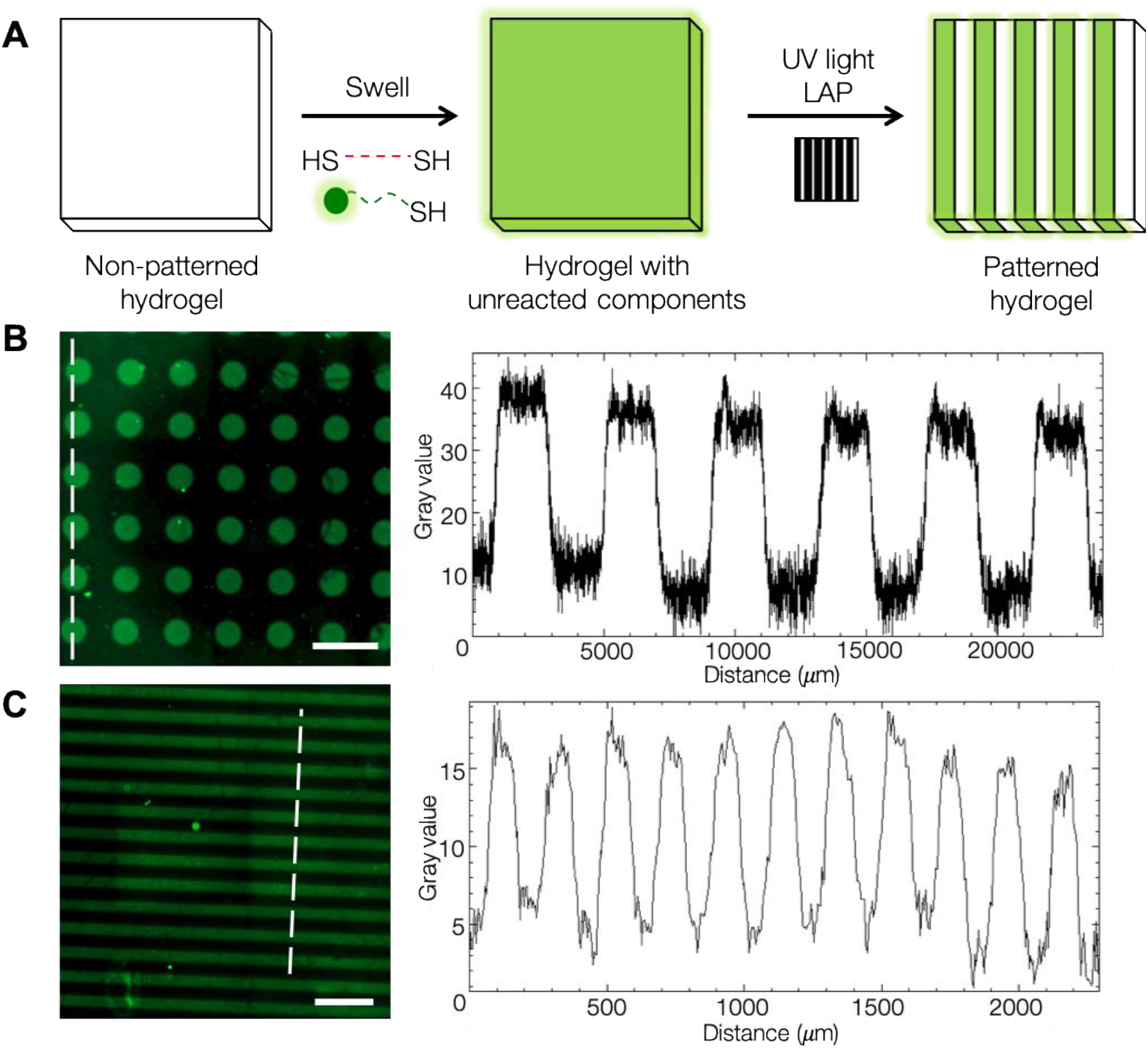
Photopatterning of hydrogels to introduce heterogeneous properties. (A) Schematic of the photopatterning process. NorHA hydrogels were swollen with thiolated molecules, covered with a photomask, and exposed to UV light, resulting in regions that underwent secondary crosslinking via light-mediated thiol-ene addition. A model thiolated fluorescent peptide was used to demonstrate patterning capabilities. Color intensity profiles showed high pattern fidelity across pattern features for (B) 200 μm diameter circles and (C) 200 μm stripe patterns; signal intensity profiles were quantified along the white dotted lines. Scale bars: 500 μm.

After establishing the photopatterning approach, we wanted to develop a patterned hydrogel model of fibrotic tissue. During the heterogeneous progression of fibrosis, the aberrant shift in healthy tissue mechanics from soft and viscoelastic to stiff and more elastic highlights the need for *in vitro* models enabling independent spatial control of both stiffness and viscoelasticity. Given the ability for multiple light-mediated thiol-ene click reactions to occur in series, our hydrogel system can model both the heterogeneity of fibrosis through photopatterning and the induction of fibrosis progression through introduction of new crosslinks to stiffen the hydrogel. Starting from an initial soft viscoelastic hydrogel, we photopatterned in additional covalent crosslinks to create stiff, less viscoelastic hydrogel regions mimicking fibrotic nodules. Atomic force microscopy (AFM) was used to characterize the elastic and viscoelastic properties of the patterned substrates. AFM nanoindentation tests demonstrated that patterned hydrogel regions undergoing secondary covalent crosslinking displayed higher Young’s moduli (16.5 ± 0.23 kPa) compared to non-patterned regions (1.78 ± 0.22 kPa) (**Figure 6A**). The viscoelasticity of the patterned hydrogel regions was characterized using a nanoindentation test followed by a dwell period in which the AFM tip was held at a constant indentation depth to measure force as a function of time. The non-patterned soft viscoelastic regions showed ~ 20% force relaxation over a period of 10 seconds, whereas the patterned stiff elastic regions showed negligible relaxation (**Figures 6B, 6C**), similar to bulk rheological measurements for homogeneous hydrogels. Importantly, this novel method for patterning viscoelasticity can be decoupled from changing stiffness. As a demonstration of this, viscoelasticity can be patterned into a stiff elastic substrate through the introduction of supramolecular crosslinks to produce regions of patterned, stiff viscoelasticity without changing the overall Young’s modulus (initial stiff elastic = 10.6 ± 0.38 kPa, patterned stiff viscoelastic = 10.7 ± 0.49 kPa) (**Figure S9**).

**Figure 6.**
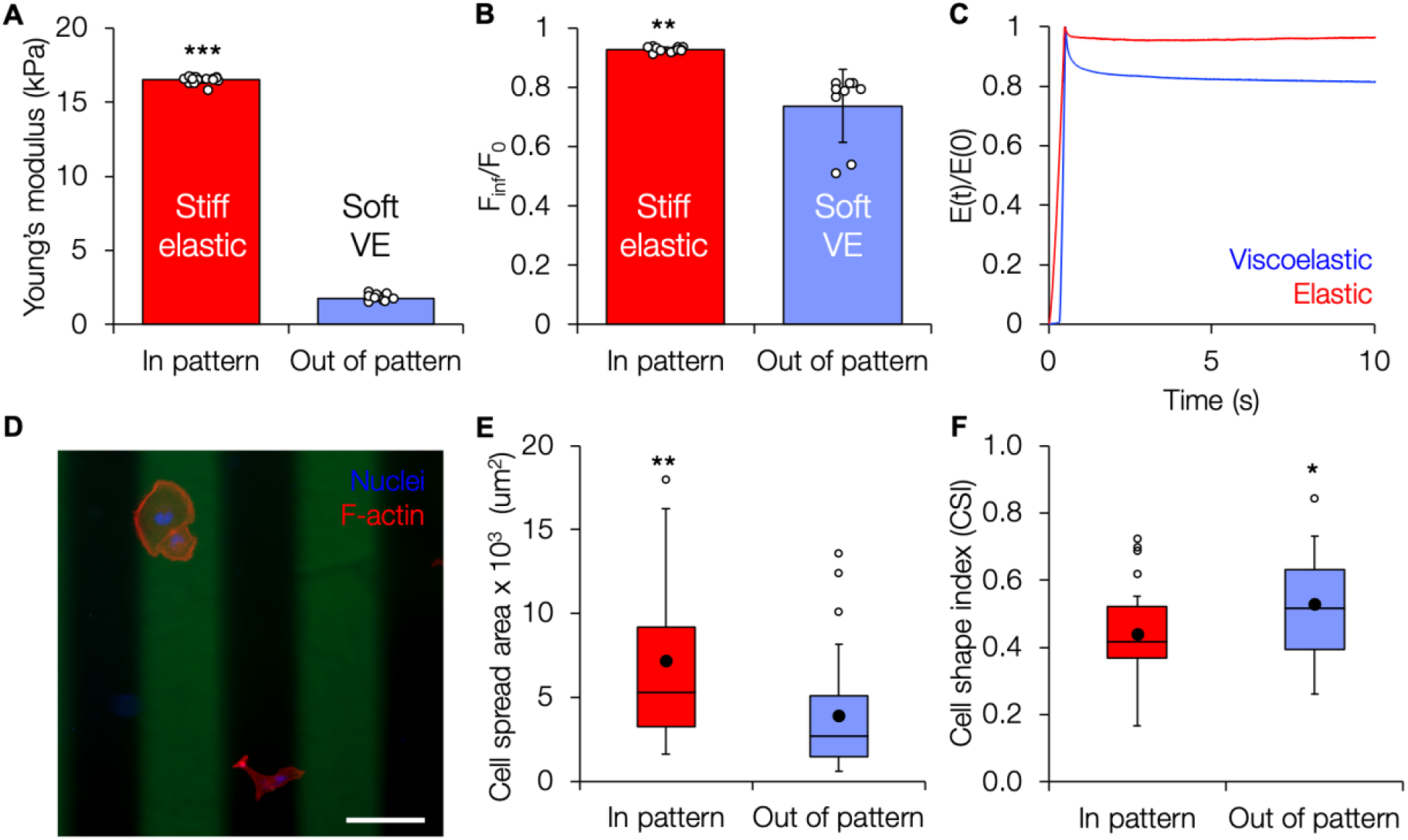
Mechanical characterization and cell response on patterned viscoelastic hydrogels. (A) Patterned (stiff elastic) and non-patterned (soft viscoelastic) regions showed differences in Young’s moduli similar to homogeneous substrates. (B) Quantification of the ratio of equilibrium force (F_inf_) to initial indentation force (F_0_) of patterned hydrogels indicates significantly greater levels of stress relaxation in the non-patterned (soft viscoelastic) regions. (C) Time-dependent Young’s modulus E(t) normalized to the initial instantaneous modulus E(0) showed ~ 20% relaxation in the non-patterned soft viscoelastic regions compared to negligible relaxation for the stiff elastic patterned regions. (D) Representative fluorescent image of LX-2 stellate cells (*red*: F-actin, *blue*: nuclei) cultured on a patterned hydrogel (200 μm wide stripes, *green*). Scale bar: 200 μm. (E) Cells on the patterned (stiff elastic) region showed significantly increased spread area compared to those in the non-patterned (soft viscoelastic) regions. (F) Cells in the patterned regions also showed significantly lower cell shape index, indicating a more elongated morphology, compared to more rounded cells in the non-patterned regions. *: *P* < 0.05, **: *P* < 0.01, ***: *P* < 0.001.

Next, we seeded LX-2 stellate cells onto soft viscoelastic hydrogels with patterned regions of stiff elastic mechanics (**Figure 6D**). Cells responded to the local mechanics of the patterned substrate and showed significantly increased spreading (**Figure 6E**) and significantly lower cell shape index (**Figure 6F**) on stiffer patterned regions. These results demonstrate the utility of this hydrogel system as a model of heterogeneous tissue mechanics.

## 4. Conclusions

This work developed an approach to make viscoelastic hydrogels via light-mediated thiol-ene addition of both covalent and supramolecular crosslinks. The use of light as a trigger for crosslinking enabled secondary modification of the hydrogel network to both increase stiffness (mimicking initiation of fibrosis) and/or modulate viscoelasticity (through the introduction of covalent and/or supramolecular crosslinks). We showed that LX-2 human hepatic stellate cells responded to the viscoelastic hydrogels by displaying reductions in spread area, MRTF-A nuclear translocation, and organization of actin stress fibers. We also used photopatterning to create hydrogels with stiff, elastic areas surrounded by soft, viscoelastic regions to mimic a heterogeneous fibrotic environment and showed that cells spread more in the stiffer patterned regions. Moving forward, we expect that this hydrogel system affording spatiotemporal control of stiffness and viscoelasticity will be useful to model a range of healthy and diseased cellular microenvironments.

## Supporting Information

^1^H NMR spectra for NorHA, CD-HDA, and CD-HA, MALDI spectra for peptides, and additional hydrogel mechanical characterization and cell analysis can be found in the supplemental file.

## Notes

The authors declare no competing financial interest.

## Acknowledgments

The authors would like to thank Dr. Tom Barker and Dr. Wei Li for providing training and use of the AFM, and Dr. Lin Han and Biao Han for providing the initial MATLAB code for AFM analysis. This work was supported by the University of Virginia and the Biotechnology Training Program NIGMS 5T32 GM008715.

