## Supplemental Figures for "Spatial control of viscoelasticity in phototunable hyaluronic acid hydrogels"

### Supplementary Figures

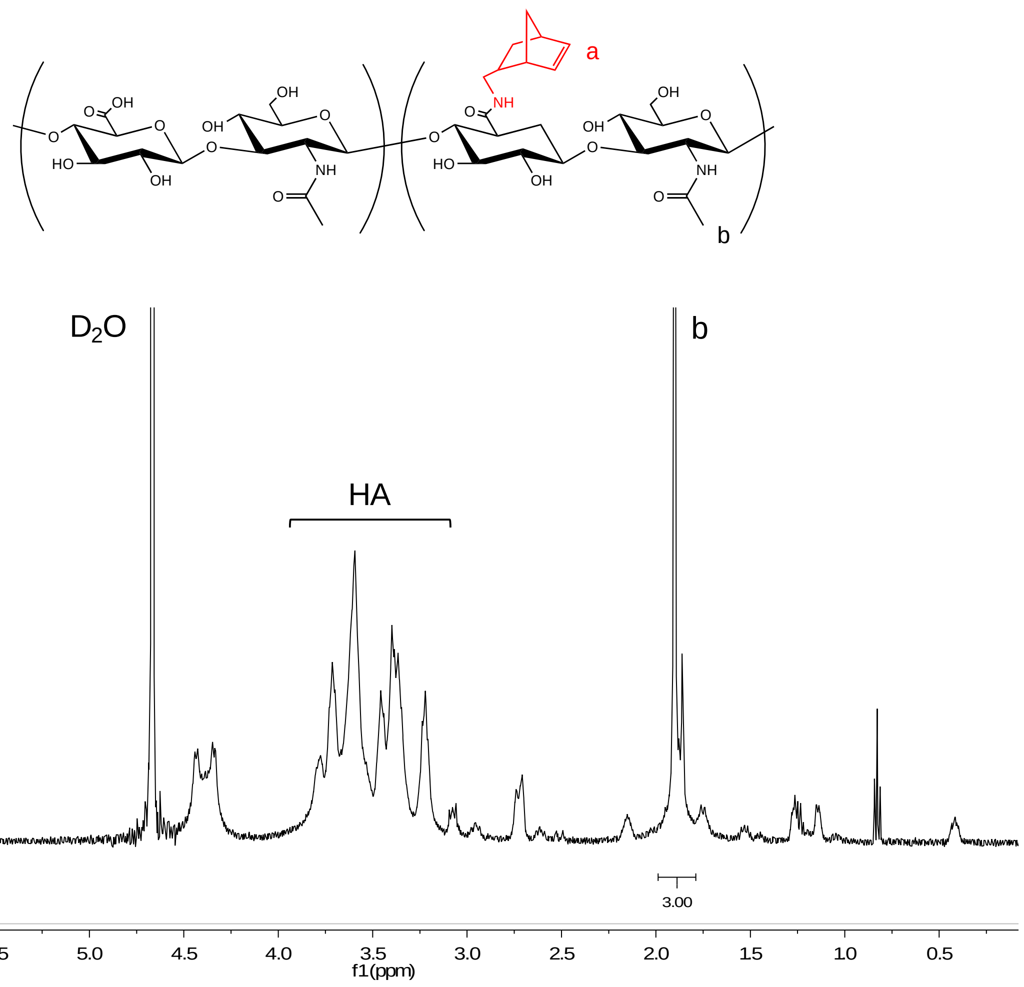


**Figure S1. $^1$H NMR spectrum of norbornene-modified hyaluronic acid (NorHA).** The degree of modification was determined to be 22%.

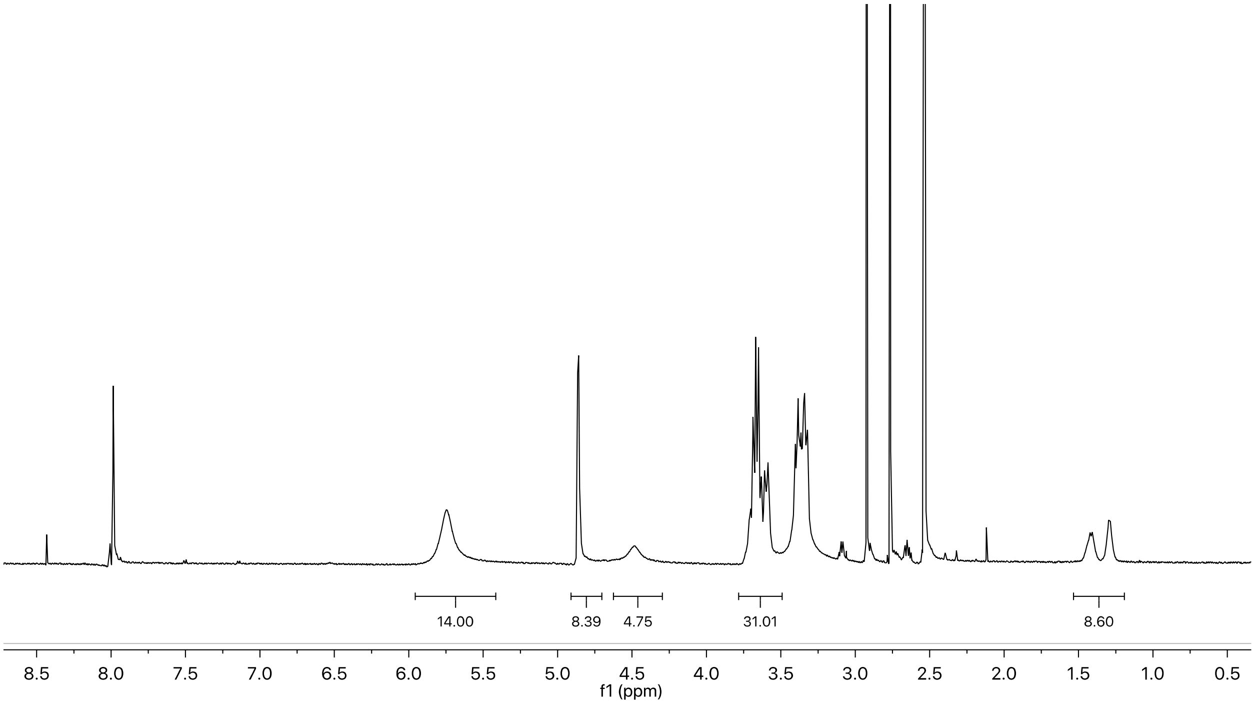


**Figure S2. $^1$H NMR spectrum of β-cyclodextrin hexamethylene diamine (β-CD-HDA).** The degree of modification was determined to be 61%.

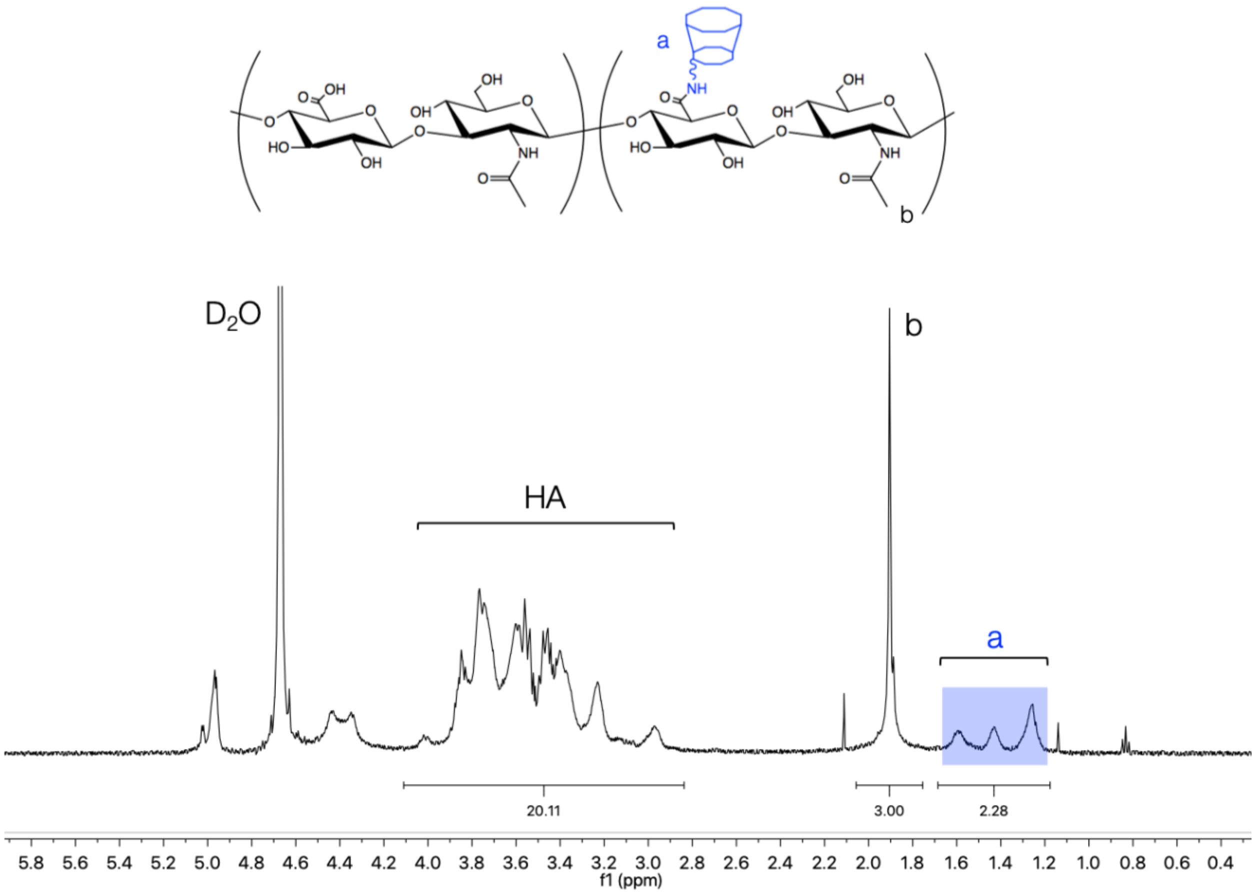


**Figure S3. $^{1}$H NMR spectrum of β-cyclodextrin-modified hyaluronic acid (CD-HA).** The degree of modification was determined to be 27%.

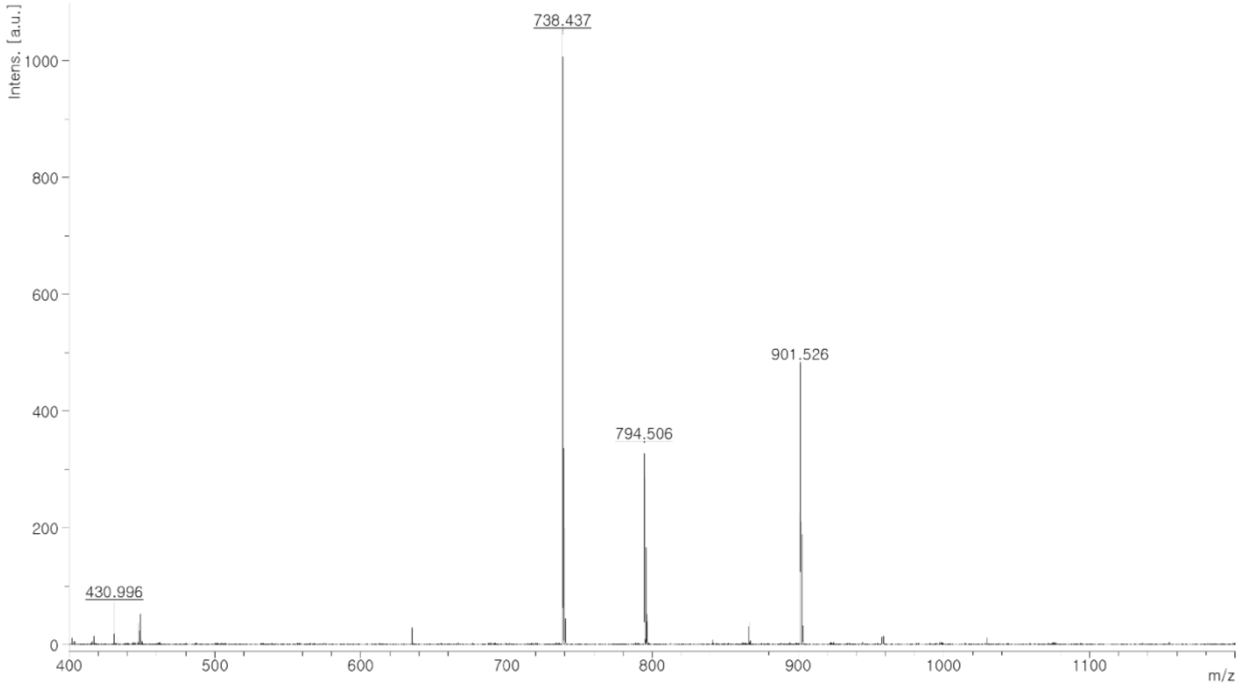


**Figure S4. MALDI spectrum of adamantane peptide with the sequence 1-adamantaneacetic acid-KKKCG.** Expected mass: 738.6 g/mol. Actual mass: 738.4 g/mol.

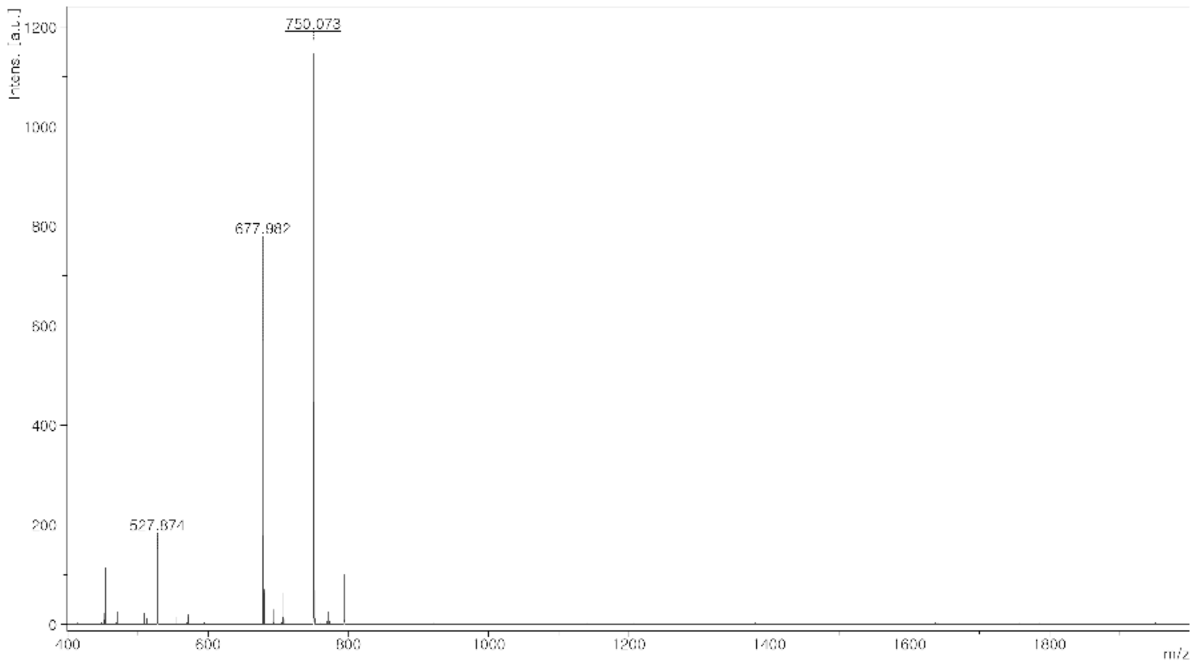


**Figure S5. MALDI spectra of fluorescent peptide.** Fluorescent peptide with the sequence Fluorescein-KKCG. Expected mass: 749 g/mol. Actual mass: 750 g/mol.

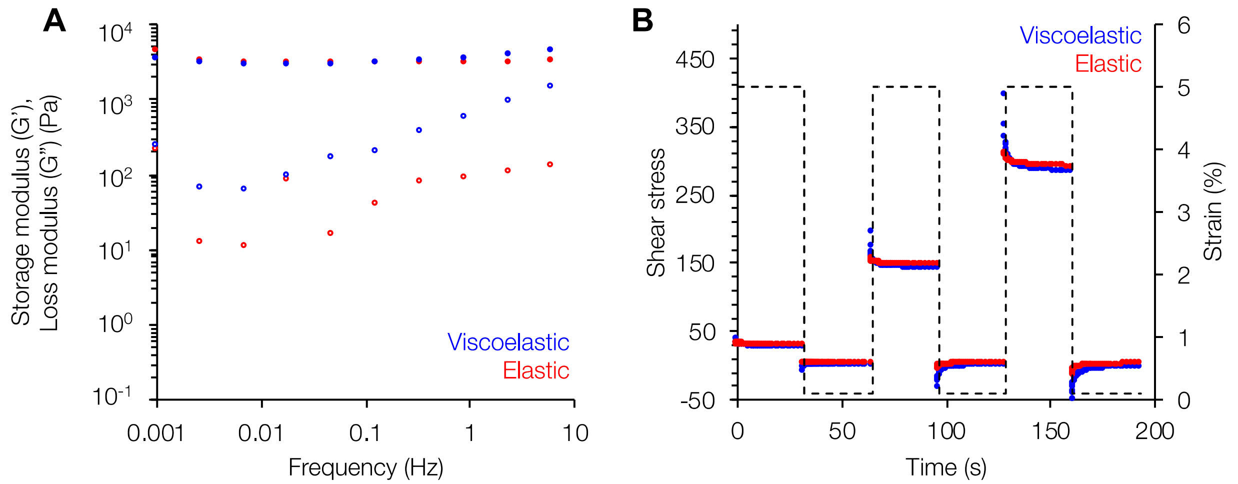


**Figure S6. Rheological characterization of stiff hydrogel groups.** (A) Viscoelastic hydrogels also showed frequency-dependent behavior with increasing loss moduli as frequency was increased, whereas the elastic hydrogel properties remained relatively constant. (B) Stress relaxation and recovery tests showed the full recovery of the mechanical properties of the viscoelastic hydrogels. For the frequency and stress relaxation tests, the stiff hydrogel groups are shown; the soft hydrogel groups can be found in Figure 2.

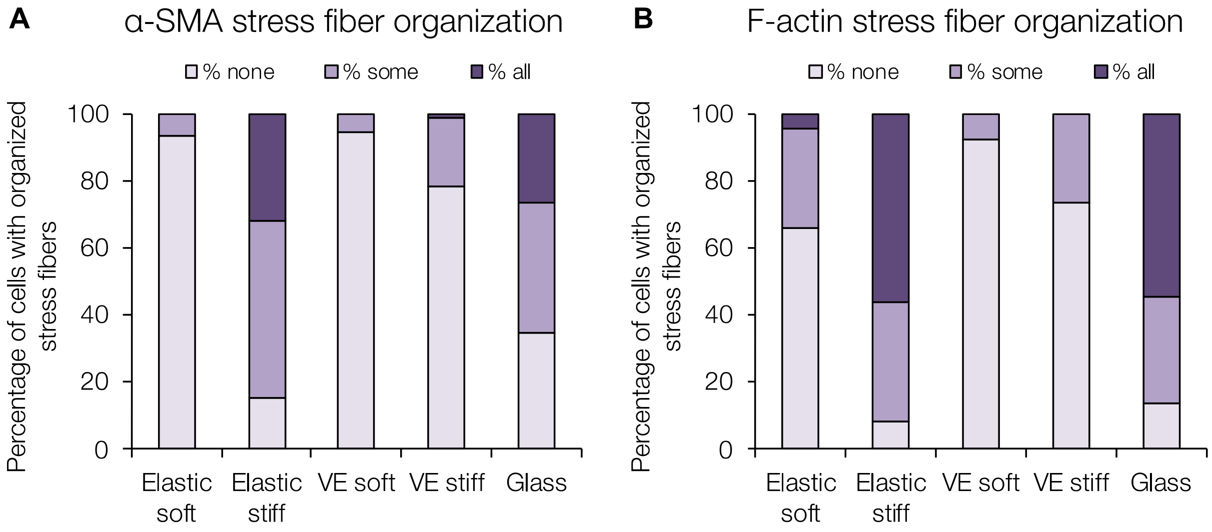


**Figure S7. Characterization of stress fiber organization.** Percentage of LX-2 hepatic stellate cells showing distinct (A) α-SMA and (B) F-actin stress fiber organization.

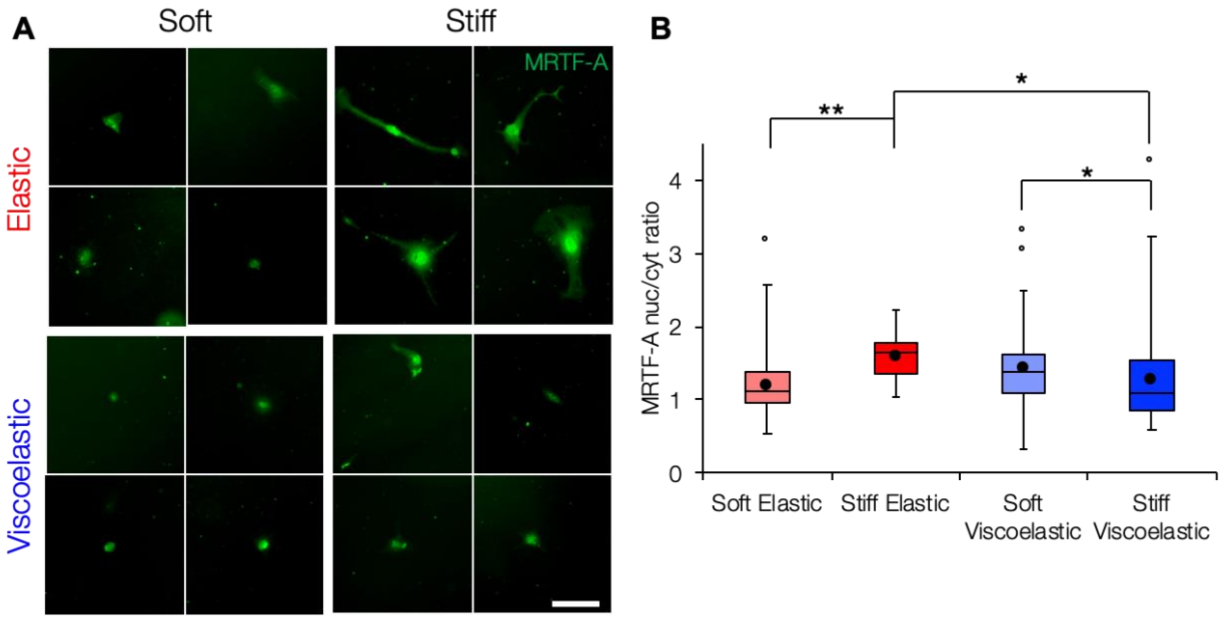


**Figure S8. Quantification of MRTF-A nuclear localization.** (A) Representative images of MRTF-A nuclear localization (*green*) in LX-2 hepatic stellate cells. (B) Quantification of the nuclear to cytosolic ratio of MRTF-A staining showed increased nuclear localization in the stiff elastic versus soft elastic groups (nuclear/cytosolic ratios of 1.61 and 1.20 respectively) while nuclear localization was similar between soft and stiff viscoelastic groups. Scale bar 100 µm. *: $P < 0.05$, **: $P < 0.01$.

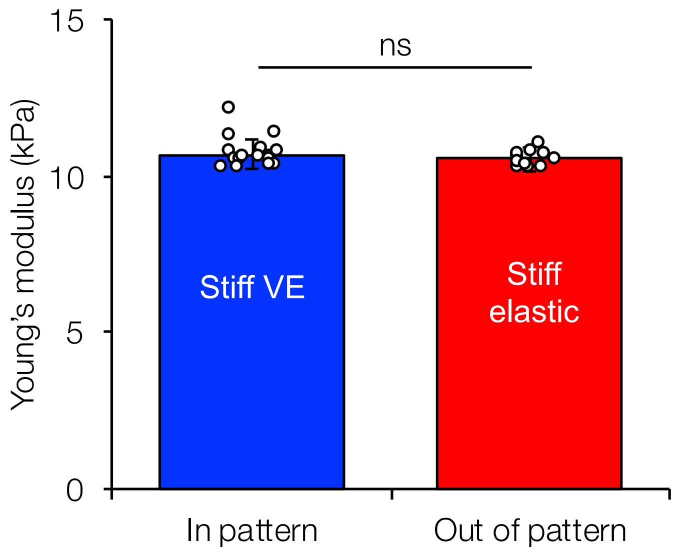


**Figure S9. Patterning viscoelasticity without changing hydrogel Young's modulus.** Atomic force microscope (AFM) mechanical characterization of patterned (stiff viscoelastic) and non-patterned (stiff elastic) regions showed comparable Young's moduli similar to homogeneous substrates.
